# A Nested Parallel Experiment Demonstrates Differences in Intensity-Dependence Between RNA-Seq and Microarrays

**DOI:** 10.1101/013342

**Authors:** David G. Robinson, Jean Wang, John D. Storey

## Abstract

Understanding the differences between microarray and RNA-Seq technologies for measuring gene expression is necessary for informed design of experiments and choice of data analysis methods. Previous comparisons have come to sometimes contradictory conclusions, which we suggest result from a lack of attention to the intensity-dependent nature of variation generated by the technologies. To examine this trend, we carried out a parallel nested experiment performed simultaneously on the two technologies that systematically split variation into four stages (treatment, biological variation, library preparation, and chip/lane noise), allowing a separation and comparison of the sources of variation in a well-controlled cellular system, *Saccharomyces cerevisiae*. With this novel dataset, we demonstrate that power and accuracy are more dependent on per-gene read depth in RNA-Seq than they are on fluorescence intensity in microarrays. However, we carried out qPCR validations which indicate that microarrays may demonstrate greater systematic bias in low-intensity genes than in RNA-seq.

## Introduction

Since the introduction of RNA sequencing (RNA-Seq) for measuring mRNA expression, one important question has been how the technology compares to microarrays in power and accuracy. Experiments have been carried out to compare microarrays and RNA-Seq, with some concluding that RNA-Seq shows a greater power, accuracy and dynamic range [1, 2] and others challenging that conclusion [3, 4].

We carried out a genome-wide gene expression experiment in a controlled setting on the yeast *Saccharomyces cerevisiae* in such a manner that the major sources of Profiling variation can be unbiasedly partitioned and quantified (Figure 1). A single extraction of mRNA from each sample was quantified by both microarrays and RNA-seq in parallel. We multiplexed each lane of RNA-seq profiling so that it exacty mirrored the eight chip per array design of the microarray platform that we utilized. This experiment allowed a direct and completely parralel investigation into the quantitative operating characteristics of RNA-seq gene expression profiling versus microarrays.

**Figure 1:**
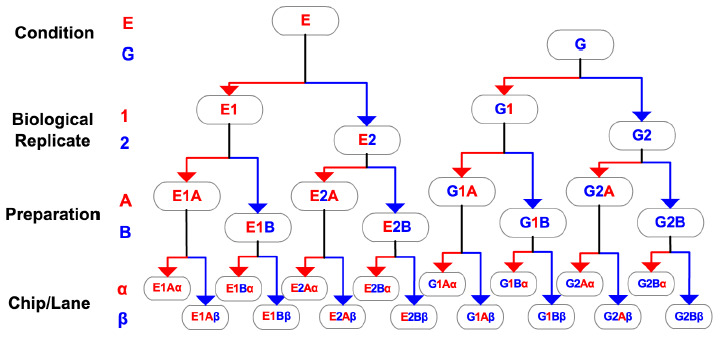
Schematic of the factorial experiment. The Condition and Biological Replicate steps were performed irrespective of technology and these materials were then utilized in a technology-specific manner (microarrays or RNA-seq) for the Preparation and Chip/Lane steps.

We use this experiment to examine the variation added by the RNA-Seq and microarray technologies. Our analysis focuses on two novel problems: decomposing the variation contributed by each technology into multiple stages, which is made possible by the nested design, and analyzing the variation as a function of gene expression intensity^1^, which is known to influence technology-specific variation in both microarrays [5, 6] and RNA-Seq [7, 8]. We find that the variance contributed by RNA-Seq is more intensity-dependent than that from microarrays, a result that is statistically significant and is robust across multiple normalization methods and *R*^2^ metrics. However, comparisons to qPCR validations show that microarray may show systematic biases in low intensities, possibly due to cross-hybridization. This has implications for the design of future experiments.

A novel characteristic of our experiment relative to previous microarray to RNA-Seq comparisons is that we utilized barcode multiplexing to combine RNA-Seq replicates on each of two sequencing lanes, meaning that technical variation added by library preparation and handling is now distinguishable from “sampling” variation added by the lane. Also notable is that our analysis takes into account the effect that intensity has on variation, which has confounded previous comparisons [9]. We thus examine which technology adds more variation as a function of per-gene depth or intensity.

## Results

### Nested Factorial Experiment

We carried out an experiment on both Agilent microarrays and Illumina RNA sequencing to investigate the effects of each source of variation on the inference of differential expression. We used the widely studied model organism *Saccharomyces cerevisiae* to investigate differential expression associated with growth in different carbon sources, glucose (G) and ethanol (E), and we introduced steps to capture three additional factors or sources of variation. The sources of variation are:

(1) **Condition**: biological condition of interest (G vs. E);
(2) **Biological Replication**: natural biological variation between clonal populations;
(3) **Preparation**: sample handling and preparation;
(4) **Chip/Lane**: technical variation associated with either technology, such as array effects or lane effects.

Factors (1)–(4) were sequentially nested, and at the stage of each nested factor, the sample was split such that each sample from the previous factor is balanced across both levels of the factor (Figure 1). Factors (1) and (2) were performed only once to produce four samples of isolated RNA, while factors (3) and (4) were technology-dependent and therefore performed in parallel with microarrays and with RNA-Seq. We took advantage of a similar design on Agilent yeast microarrays (8 hybridizations per chip) and Illumina RNA-Seq (8 indexed samples per lane) to mimic the same approach across the two technologies, resulting in 16 microarray and 16 RNA-Seq profiles that show the amount of variation added at each stage. The RNA-Seq experiment achieved a depth of 170.8 million reads, with depths of 89.6 million and 81.1 million on each of the two lanes.

### Differential Expression

The biological goal of this experiment is to infer differential gene expression in *Saccharomyces cerevisiae* strain DBY12000 (S288c Hap1+ Mat *a*) cultivated in balanced growth conditions in chemostats using either glucose or ethanol as the sole carbon source (condition of interest). The chemostat device helped minimize any variations in growth conditions (such as physiological state, temperature, nutrient composition, etc.) so the study could directly interrogate the factor of interest: transcriptional responses to different carbon sources [10]. We used a linear model with t-statistic shrinkage to detect differential expression in each case, after log-transforming the RNA-Seq counts and fitting precision weights to each observation [11, 12]. We specifically tracked a biologically relevant set of 30 genes known to be involved in processes relevant to glucose or ethanol metabolism, namely gluconeogenesis, glycolysis, the TCA cycle and the pyruvate branchpoint [13].

One goal is to assess to what extent RNA-Seq and microarray experiments agree in their assessment of differential expression. Figure 2a compares the estimated log_2_(G/E) fold change ratios between the two technologies, with the opacity of each point determined by the quantile of the microarray intensity or RNA-Seq read depth, whichever is lower. The microarray intensity of each gene was calculated as the average cy5 flourescence intensity across all samples (where cy5 is the channel corresponding to the samples of interest), while RNA-Seq depth was calculated as the total number of reads mapping to the gene across all samples. We also show 30 biologically relevant genes highlighted in color. Overall the two technologies show a correlation of 0.855, but this correlation is highly dependent on read depth and microarray intensity, with the lowest-intensity genes exhibiting the greatest noise and therefore the lowest correlation.

**Figure 2:**
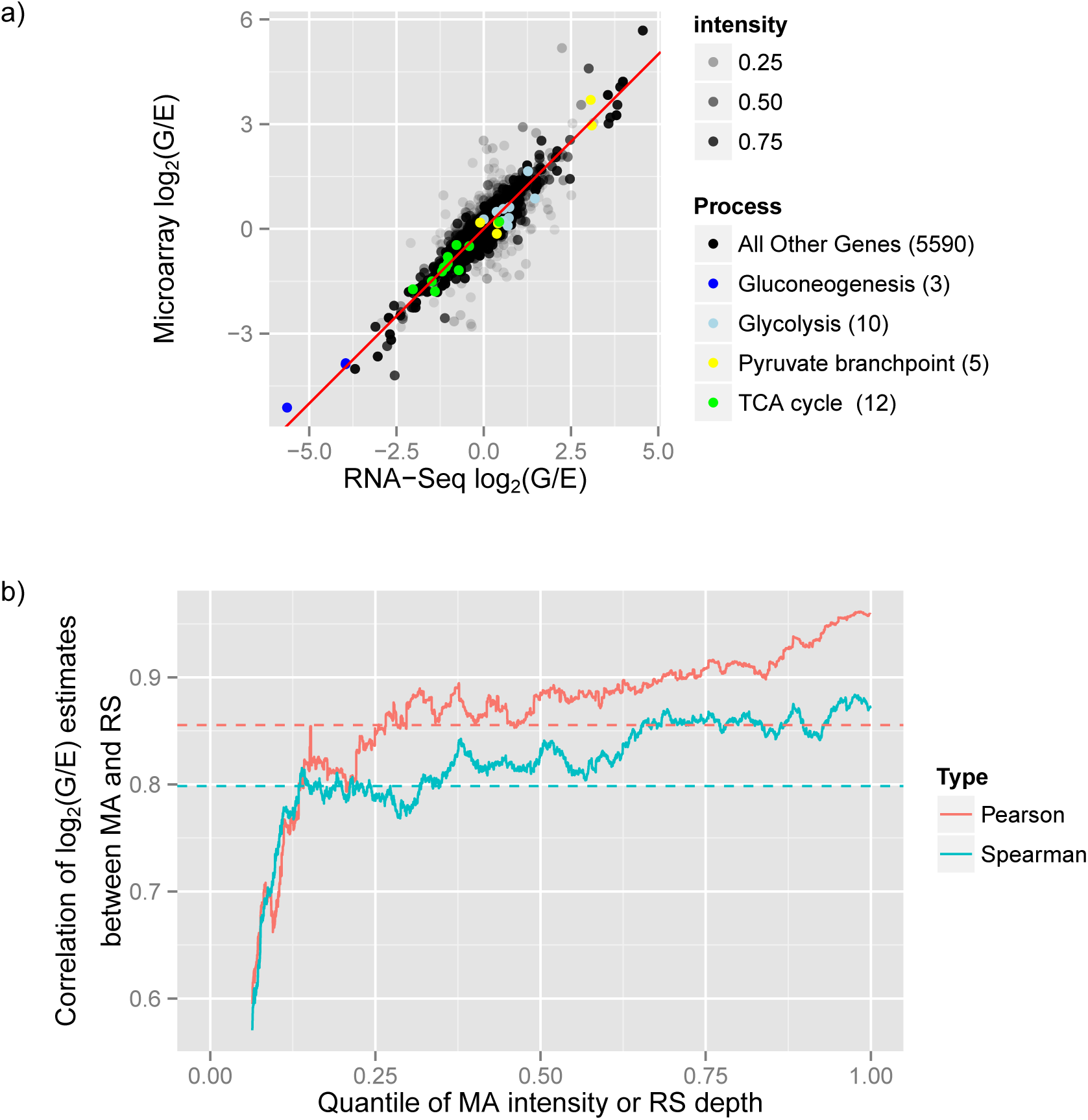
a) Comparison between the log_2_(G/E) (log fold-change) estimates measured with RNA-Seq or with the microarray. The transparency of the points corresponds to the quantile of the intensity in microarray or RNA-Seq, whichever is lower. 30 genes from biologically relevant pathways are highlighted in color. b) Pearson or Spearman correlation of log_2_(G/E) estimate between microarray and RNA-Seq, within a 500 gene window rolling over microarray/RNA-Seq intensity ranking. The correlation of all genes (0.855 Pearson, 0.798 Spearman) is shown as a horizontal dashed line.

Figure 2b shows how the Pearson and Spearman correlations between microarray and RNA-Seq effect size estimates depend on per-gene intensity, using a rolling window of 500 genes, ordered by intensity (again determined by the quantile of the microarray intensity or RNA-Seq read depth, whichever is lower). The Pearson correlation varies from 0.595 to 0.961, while the Spearman ranges from 0.57 to 0.884. The 30 genes in our biologically relevant set showed a 0.985 correlation of effect size estimates, which is understandable since almost all lie in the top 10% of read depth and microarray intensity. The microarray and RNA-Seq assays identified as significantly differentially expressed 28 of the 30 biologically relevant genes at estimated FDR ≤ 5% [14], and the two technologies agreed on the direction of the change for all of these genes. This suggests that there is little difference between the two technologies in terms of estimating differential expression of high-intensity genes.

### Percentage of Variation Explained

The primary goal of the factorial experiment is to determine the relative amount of variation added at each stage of the experiment for each of the two technologies. Based on the correlation matrix (Figure S1), the RNA-Seq and microarray assays easily distinguished between the ethanol and glucose samples, and showed clustering within the chip/lane replicates, as expected from the experimental design. Any analysis should, however, consider the intensity-dependence of the variation that each technology contributes. We calculated the proportion of variation explained by the Condition, Biological, and Preparation factors, as well as the residual variation due to the chip or lane, using a nested ANOVA analysis (**Materials and Methods**). We computed this breakdown separately for each gene and smoothed the result across microarray intensity RNA-Seq depth using the local regression method, LOESS (Figure 3).

**Figure 3:**
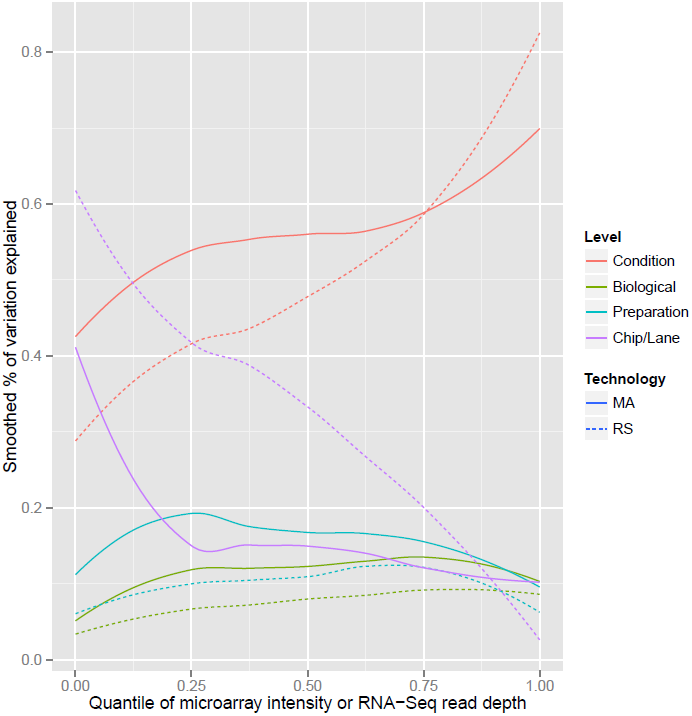
Percent of variance explained by each nested level of the experiment as computed by an ANOVA adjusted *R*^2^, smoothed using LOESS across the intensity quantiles.

In both technologies at almost all intensities, the largest sources of variation were the treatment (ethanol versus glucose) and the chip/lane, in a tradeoff that depended strongly on intensity or read depth. The analysis indicated that the variance due to RNA-Seq lane at low-intensity genes was greater than that due to microarray chip. This difference is statistically significant: when the genes are divided into 10 bins based on depth or intensity, all bins in the 80% lowest intensity show a highly significant difference between the amount of variation added by the microarray chip versus the sequencing lane (Figure S2). These conclusions were robust across multiple methods of microarray and RNA-Seq normalization (Figure S3). To ensure that this discrepancy did not arise from the difference between the discrete count data from RNA-Seq and the quasi-continuous fluroescence intensity data from the microarray, we also calculated an alternative *R*^2^ designed for count data to determine the proportions of variance explained (**Materials and Methods**) [15], and observed almost no difference (Figure S4).

To examine the variation added by the chip/lane level more directly, we also measured the difference in log fold change estimate when the same sample was run on two chips or two lanes. We computed the Pearson and Spearman correlation of log fold changes between chips or between lanes in overlapping windows of 500 genes ordered by intensity as above (Figure 4a), as well as a LOESS-smoothed curve of the absolute value of the difference (Figure 4b). Both analyses confirm that the difference between RNA-Seq lanes is more intensity-dependent than the difference between microarray chips, with a particularly great disagreement in low-intensity genes. One notable question is whether this effect can be mitigated by effect size shrinkage, such as the DESeq2 software, which is designed to improve the stability of estimates for low-depth genes [16]. Figure 4 shows that DESeq2 causes the absolute difference in log fold changes to decrease, but does not substantially improve the correlation in any but the lowest-intensity windows. This suggests that effect size shrinkage can make estimates less variable between lanes, but does not remove the intensity dependence.

**Figure 4:**
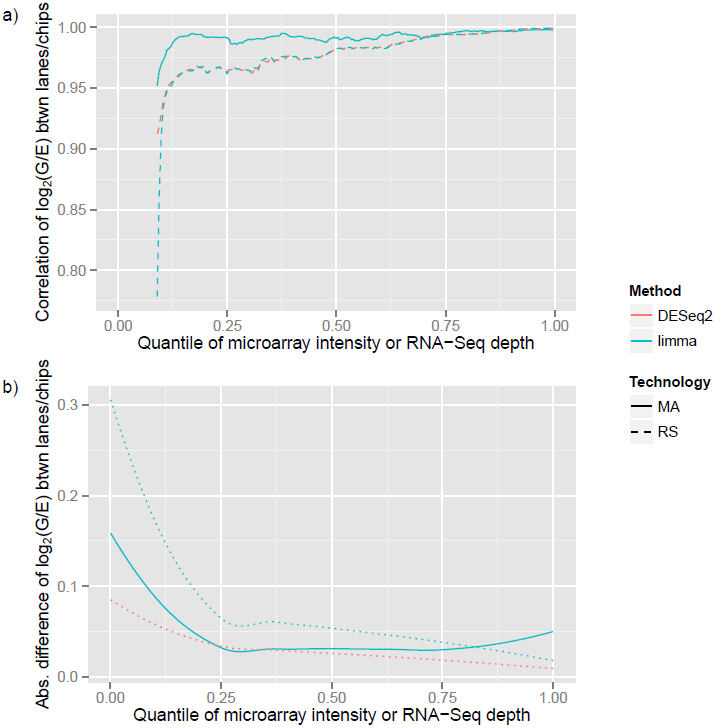
Comparisons of the log_2_(G/E) estimates within each technology, comparing the estimates computed for each of the two chips in a microarray or two lanes in RNA-Seq, using the same library preparations. a) Pearson correlation between the two lanes of RNA-Seq or two chips of microarrays, within a 500 gene window rolling over intensity. b) Absolute value of the log_2_(G/E) estimate difference between chips in microarray or lanes in RNA-Seq, smoothed using LOESS.

### Validation of Low-Intensity Genes

These results do not necessarily indicate that microarrays are more accurate at low intensities than RNA-Seq, only that they show greater consistency between replicates. Each technology may still possess biases that cause their measurements not to reflect the underlying mRNA abundance levels. To examine the accuracy of each technology more directly, we chose 13 genes in the bottom 20% of intensity for both technologies, for which the estimates of the log_2_(G/E) fold change disagreed by at least 1.0 between the technologies. On these low-intensity contested genes, we performed quantitative PCR (qPCR) on the same mRNA as the experiment to provide an independent validation (**Materials and Methods**). As shown in Figure 5, the qPCR results agreed much more closely with the RNA-Seq estimates than with the microarray: the correlation of estimated RNA-Seq and qPCR log fold changes is 0.671 (p-value ≈ 0.01), while the microarray to qPCR correlation is 0.016 (p-value ≈ 1).

**Figure 5:**
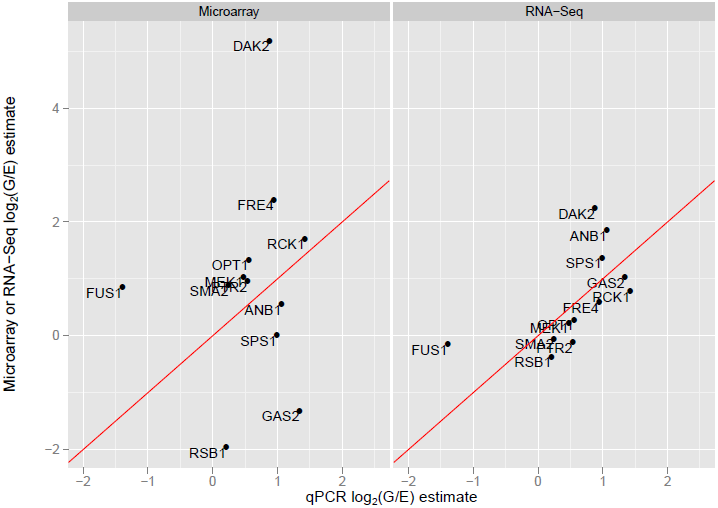
Comparison of the log fold change estimates measured with qPCR compared to estimates from the microarray or RNA-Seq, for 13 selected low-intensity genes that disagreed between RNA-Seq and microarray.

This leads to the question of whether the disagreement between microarrays and qPCR results from greater variation present at low-intensity, or rather systematic bias introduced by the technology, such as cross-hybridization or fold-change compression. Towards that end, we examined the distribution of normalized measurements across all biological, technical and chip/lane replicates from the 6 genes for which microarrays most strongly disagreed with qPCR (Figure 6). Even though these genes are low-intensity, in most cases the variation within microarray and RNA-Seq measurements was very small compared to the disagreement between the technologies. In a dramatic example, GAS2 shows a reversed fold change relative to qPCR and RNA-Seq, but shows very little within-condition variation in any technology. Of these 6 genes, only OSW1 could plausibly be caused by fold change compression, since in other cases the microarray effect was greater than or a reversal from the RNA-Seq and qPCR observations [17]. This suggests that the issue within these genes was not random variation present in low-intensity genes or fold change compression, but rather a bias that led to a spurious but statistically significant effect size estimate.

**Figure 6:**
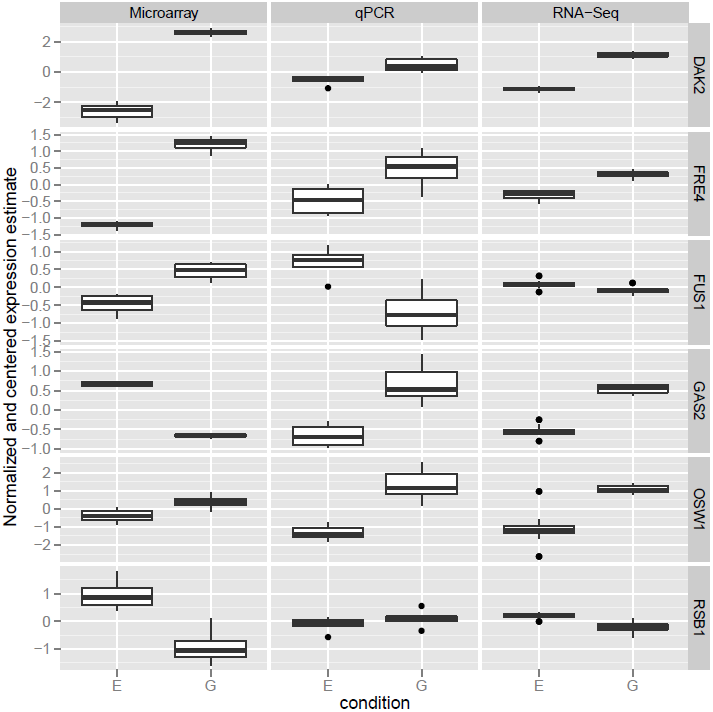
Boxplots comparing the normalized and centered expression values of microarrays, RNA-Seq, and qPCR of the 6 genes for which microarray and qPCR most disagreed. This shows that in many cases, microarray measurements were very consistent between biological, technical and chip replicates. This suggests that the problem is not variation at low-intensity microarray measurements, but rather bias.

### Gene Set Enrichment

Many gene expression analyses seek to determine sets of genes whose expression changes within a particular condition in order to draw biologically relevant conclusions [18, 19]. We thus looked for enriched sets of genes within each technology using a Wilcoxon rank-sum test comparing genes within a gene set to those outside it [20, 21].

Many enrichment tests look for genes that are unusually high or low on a ranked list, often ranked by statistical significance. However, ranking by statistical significance has been noted to lead to a confounding effect in RNA-Seq, where highly-expressed or high-depth gene sets are spuriously marked as significant [22, 23]. An alternative is to perform enrichment analysis on log fold change estimates, which would be expected to be less confounded with depth [24]. To demonstrate this effect in our data, we performed gene set enrichment using either p-values or fold change estimates from each technology, then assigned the gene sets into 5 bins based on each gene set’s median intensity (Figure S5). We see that the enrichment p-value histogram is highly conservative for low-intensity gene sets when differential expression p-values are used, and is less influenced by intensity when the log fold change estimate is used instead. Notably, even though we found the per-gene variation to be more intensity dependent in RNA-Seq than in microarray, the intensity dependence of gene sets is similar between the two technologies. Another reason to use the fold-change estimate is that the significance metric is more technology dependent: the p-values for differential expression show only a moderate Spearman correlation of 0.544 between microarrays and RNA-Seq, while the fold change estimate shows a higher correlation of 0.798. We thus chose to use the estimate of effect size rather than statistical significance to evaluate gene set enrichment.

The results of our gene set analysis of both the RNA-Seq and the microarray data are shown in Table S2, and the distributions of the log fold change of some of the most significantly enriched gene sets are shown in Figures S6–S8. The enrichment of gene sets for carbohydrate catabolic process and glycolysis in glucose and of mitochondrial respiratory chain and ATP synthesis coupled proton transport serve as confirmation that the experiment captures the difference between the two metabolic states. Two of the three most significantly enriched sets are cytoplasmic translation and mitochondrial translation, for which expression is higher in glucose and in ethanol, respectively. As the rate of respiration in the mitochondria is higher in ethanol than in glucose, this suggests that increase in mitochondrial activity is reflected in a tradeoff of translation from the cytoplasmic ribosomes to the mitochondria. Another notable result is that genes in iron ion homeostasis and ferric-chelate reductase activity are higher expressed in ethanol than in glucose. This is likely due to the important role of iron transport and reduction in heme biosynthesis, which in turn is necessary for the electron transport chain and other respiratory activity [25, 26, 27].

We identified higher expression of cytokinesis and cellular budding genes in ethanol, even though growth rate was kept equal between the two samples. Mitochondrial inheritance and distribution is known to be actively regulated by the budding tip and to be necessary for equal and efficient distribution of the organelles [28, 29], and indeed some differentially expressed genes in our experiment, such as UTH1, have been identified as being related to both cell wall biogenesis and mitochondrial division [30, 31, 32]. Our results suggests that this process of mitochondrial inheritance may be transcriptionally regulated in response to the metabolic state or level of respiratory activity.

## Discussion

We demonstrate an experimental and statistical approach for determining the variation added at each stage of a microarray or RNA-Seq experiment. We determined that RNA-Seq shows a greater degree of intensity-dependent variation than do microarrays, with particularly high variance for low-intensity genes, and that the intensity-dependent component was contributed mostly by the chip or lane level. With qPCR validation, however, we discovered that microarrays appear to possess some systematic biases in their estimation of differential expression for low-intensity genes. Since this bias appears to be consistent across biological, technical and chip replicates, it likely cannot be solved or even detected by performing additional replicates on the microarray platform.

Our results have implications for the design of microarray and RNA-Seq experiments meant to identify differential expression. While other experiments will vary in the amount of variation added at the biological stages, that variation is likely to be intensity-independent as it was in our study, meaning our qualitative conclusions are likely to hold. For high-intensity genes there is little difference either in the genes called significant or the estimate of effect size between RNA-Seq and microarray, and therefore the decision of which technology can be made on other criteria, such as cost. However, in low-intensity genes, the RNA-Seq technology tends to add greater variation, leading to lower statistical power and greater uncertainty in expression estimates. Microarrays, while more consistent in their estimates across technical replicates, show systematic biases that confound differential expression detection. This suggests that studies for which low-expressed genes are of special interest should be performed cross-platform.

More importantly, our study has demonstrated that the intensity-dependent nature of variation must be taken into account in future technology comparisons and quality control experiments. Our approaches of dividing genes into intensity bins, observing correlations within overlapping windows, and smoothing per-gene values using LOESS showed trends and differences in the technologies that would have been missed using aggregate statistics. Finally, we have demonstrated how our experimental data is an appropriate benchmark for comparing statistical analysis methods and for developing experimental recommendations, as it analyzes a well-studied system, includes variation at each stage of an experiment, and compares RNA-Seq and microarrays directly on the same biological samples. We expect future research by ourselves and others will extend our conclusions and develop them further.

## Materials and Methods

### Growth of Yeast in Chemostats

Yeast haploid strain DBY 12000 FY, a S288C derivative containing the wild-type HAP1 allele, was used in this experiment. A single colony was split into 2 overnight cultures, one containing ethanol limitation medium with 60 C-mM (for detailed formula see http://www.princeton.edu/genomics/botstein/protocols/Ethanol-30.html) and the other contains glucose limitation medium with 27 C-mM Carbon (see http://www.princeton.edu/genomics/botstein/protocols/Dex_lim.htm) as the carbon source. 1 mL of each culture was used to inoculate a chemostat (see http://www.princeton.edu/genomics/botstein/protocols/BotsteinLabChemostatSetUpmodpH.pdf). After the chemostats reached stasis, 10 ml culture was collected and frozen at −80C. A second set of biological samples were prepared the same way on a different day.

### Gene Expression Profiling

RNA was harvested from each of the 4 samples (see http://dunham.gs.washington.edu/MDchemostat.pdf). The 4 RNA samples were then processed twice on different days using the Agilent Quick Amp Labeling kit (Part no. 5190-0424) to produce 8 cRNA libraries, each of which was then hybridized on two separate chips (Yeast Expression 8x15K arrays) on different days according to the factorial design. After the arrays are washed and scanned, features are extracted using the Agilent Feature Extraction software to determine red and green intensities.

The same 4 RNA samples were also processed using the Illumina TruSeq RNA Sample Prep v2 LS protocol. Each sample was prepared twice up to the 3’ end adenylation, then each of the 8 preparations was split into two aliquots, after which each was indexed and amplified to complete the RNA-Seq library preparation. The two groups of 8 samples that were indexed together were each mixed at equimolar concentrations and sequenced on separate lanes on the same Illumina HiSeq 2500 flowcell, to produce 141bp reads. In total, we produced profiles from 16 RNA-Seq samples and 16 microarray samples with identically nested experimental designs.

### Quantitative PCR

To perform quantitative PCR, the four RNA samples were treated with RNase-Free DNase Set (Cat # 79254, Qiagen) column digestion to remove DNA in the samples. Total RNA was quantified using Quant-iT RNA assay Kit (Q33140, Invirogen) and Biotek Synergy Mx plate reader. 750 ng total RNA was used in the first strand cDNA synthesis (SuperScript III Reverse Transcriptase, Cat # 18080, Invitrogen). Prevalidated FAM-MGB Taqman probes and primers mix for all candidate genes were ordered from Life Technologies. 96 well plates (Cat# N8010560, ABI) and optical adhesive cover starter kit (Cat # 4313663, ABI) were used. qPCR reactions were set up by combining Taqman Gene Expression Master Mix (Cat # 4369016, Life Technologies) and individual probe plus primer mix. 20 l qPCR reaction was run on ABI 7900 HT Sequence Detection System using the following thermal protocol: 50C, 2 min; 95C, 10 min; 40 cycles of 95C, 15 sec and 60C, 1 min; 95C, 15 sec; 60C, 15 sec; 95C, 15 sec.

Any candidate genes with more than one band on agarose gel after qPCR were excluded from further analysis. Three replicates of RT-qPCR for each RNA sample starting from cDNA synthesis to qPCR reaction were performed. Each measurement was normalized based on the average across all genes in a biological replicate, and the log-fold change was estimated based on the difference in average number of cycles between ethanol and glucose samples.

### Preprocessing and Statistical Analysis

We used `bowtie2` with the default set of parameters to map the RNA-Seq reads to the yeast genome, and used `htseq-count` to match reads against the *S. cerevisiae* R64 release of the reference genome from the Sacchromyces Genome Database, available from http://downloads.yeastgenome.org/sequence/S288C_reference/genome_releases/. Microarrays were normalized after averaging all preparation and chip replicates within each biological replicate to create two E replicates and two G replicates. The matrix of RNA-Seq counts was first pooled within preparation and chip replicates, then was transformed using `voom` from `limma`, which also computed precision weights for each observation [12]. This pooling was necessary for differential expression because while the replicates at later nested stages introduced variation, they were not full biological replicates, and treating them as replicates for limma would be pseudoreplication that underestimates the within-group variation [33]. We tested for differential expression in each technology using a linear model with empirical Bayes shrinkage of T-statistics, implemented by `limma` version 3.20.9 [11].

### Estimating Contributions to Variation

For calculations of the percent of variation explained, the microarray log fold changes and the RNA-Seq log-transformed counts were compared, with the counts first transformed using `voom` [12]. We compared multiple methods of normalizing both the microarray and RNA-Seq data between samples, using implementations in the `normalizeBetweenArrays` function in `limma` as well as the TMM [34] and RLE [35] methods for RNA-Seq, but none made a qualitative difference in the resulting conclusions (Figure S3).

For each gene, in both the normalized RNA-Seq data or the microarray fold-change values, we calculated the adjusted-*R*^2^ based on three linear models, which differed in which levels of the experiment were parameterized and which were left as residual variation. The three models were treatment, treament × biological, and treatment × biological × preparation. Each takes the form

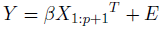

where

- *p* is the number of degrees of freedom used by the parameters in the model: respectively 1, 3, and 7 for the three models
- *g* is the number of genes
- *Y* is the *g* × 16 matrix of red/green fold changes in microarray data or the log-transformed counts of RNA-Seq data
- *X*_1:*p*+1_ is the first *p*+1 columns of the experimental design matrix (Figure S9), which includes the intercept and the *p* terms fit in the model
- *β* is the *g* ×(*p* + 1) matrix of terms
- *E* is the *g* × 16 matrix of normally distributed residuals

An adjusted *R*^2^ was found for each of the models 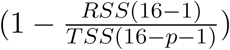
, as shown:

| Model | $p$ | Adjusted $R^2$ |
| --- | --- | --- |
| Treatment | 1 | $R_T^2$ |
| Treatment×Biological | 3 | $R_B^2$ |
| Treatment×Biological×Preparation | 7 | $R_P^2$ |

The percent of variation explained by each level was reported for each gene as 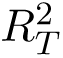 for the treatment variation, 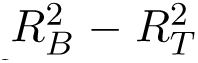 for the biological variation, 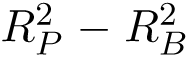 for the preparation variation, and 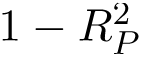 for the residual variation that occurs on the chip or the lane. This metric is meant to break down the amount that the quantification of expression, and therefore the differential expression estimation, is affected by each level. The *R*^2^ estimates were then smoothed using LOESS across all genes based on their microarray intensity or RNA-Seq read depth.

Since RNA-Seq data consists of counts rather than continuous measurements, we considered the possibility that the linear model, and therefore the *R*^2^ measure, might not capture the distribution of variation correctly. We thus also calculated two alternative *R*^2^ generalized across exponential families: 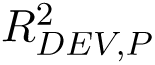 which assumes a Poisson model for the residual noise, and 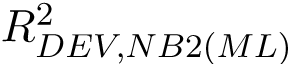, which assumes a negative binomial model [15]. This metric is defined as the decrease in Kullback-Leibler divergence between the observations and the expectation under the full model versus the divergence between the observations and the expectation under a null model. The standard *R*^2^ used in linear models with normal residuals can be viewed as a special case of this metric, so this divergence is a useful choice for comparing the count values to the traditional *R*^2^ used in the microarray [36].

We used the edgeR package to perform a negative binomial fit on each gene and estimate the dispersion parameter. We define *y_i,j_* as the unnormalized count of gene *i* in sample 
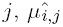
 as the expected value of *y_i,j_* under the full model (using `fitted` on an edgeR fit), *ȳ_i, j_* as the expected value using only an intercept term and the library size normalization, 
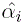
 is the dispersion of gene *i* estimated using `edgeR`, *n* is the number of samples (16) and *p* is the number of parameters estimated by the model (1 for the condition effect, 3 for condition × biological, and 7 for condition × biological × preparation). The alternative *R*^2^ estimates for the Poisson and negative binomial models can then be calculated as

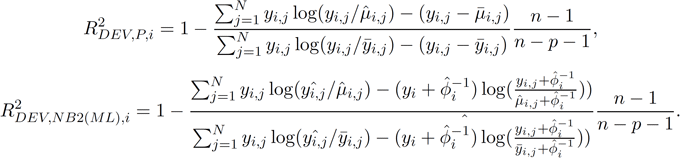

The same three models fit in the traditional linear model were fit using Poisson or negative binomial regression, using edgeR’s implementation:

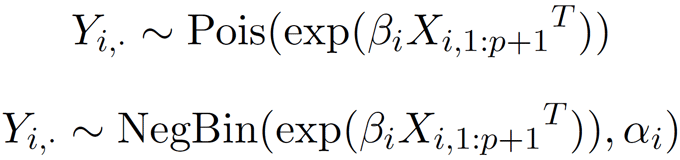

then their respective 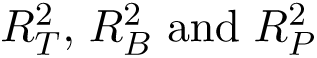 were calculated for each gene based on the difference in adjusted alternative *R*^2^ between each model. Figure S4 compares the Poisson and negative binomial models to the traditional *R*^2^ on log TMM-normalized counts. TMM was chosen since it is the default used for edgeR normalization, and Figure S3 shows that TMM produces similar results to other normalization methods.

## Computer Code and Data

Reproducible R code and the data can be obtained from https://github.com/StoreyLab/factorial-MA-RS.

## Footnotes

1 Throughout this paper, we utilize “intensity” of a gene as a term for both fluorescence intensity in microarrays and per-gene read depth in RNA-seq, both of which are measurements of a gene’s abundance subject to technology-specific biases and sources of variation.

## Supplementary Figures

**Figure S1:**
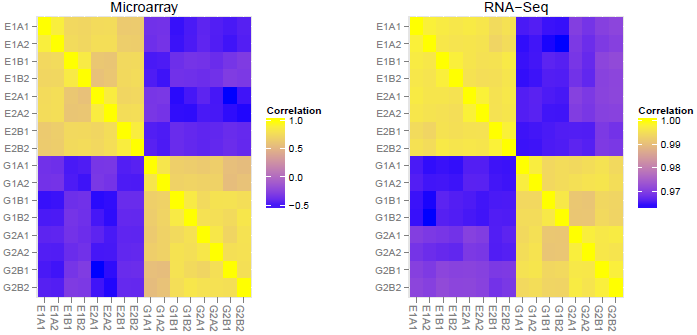
Spearman correlation matrices of microarray and RNA-Seq experiments. The microarray correlation matrix uses the log(cy5/cy3) fold change, which means the correlation can be negative, while the RNA-Seq matrix uses the raw counts, which means the correlation is always close to 1.

**Figure S2:**
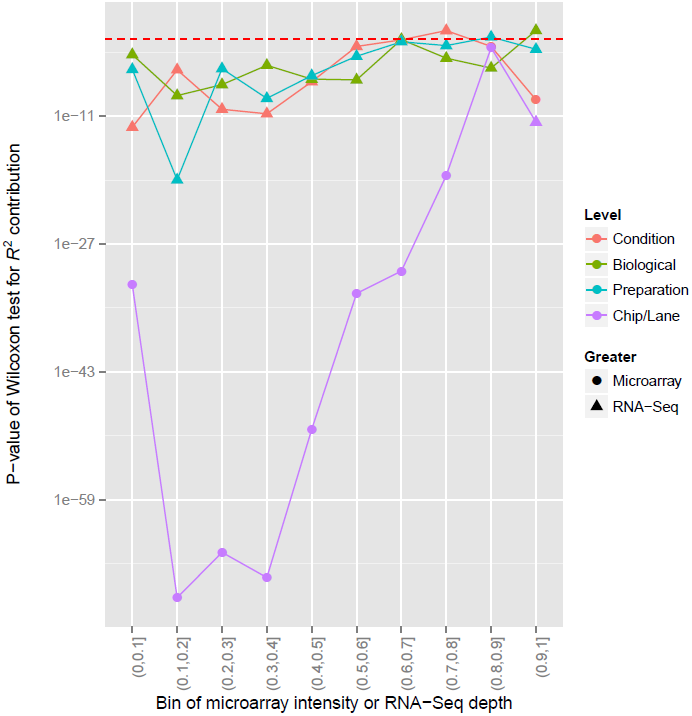
Wilcoxon rank-sum test p-values comparing the percent of variance explained at each level between RNA-Seq and microarrays, in each of 10 bins based on the intensity quantile. The shape of each point indicates whether RNA-Seq or microarrays showed higher *R*^2^ at that stage. The p-values are shown on a log scale, with p = 0.05 shown as a horizontal dashed line.

**Figure S3:**
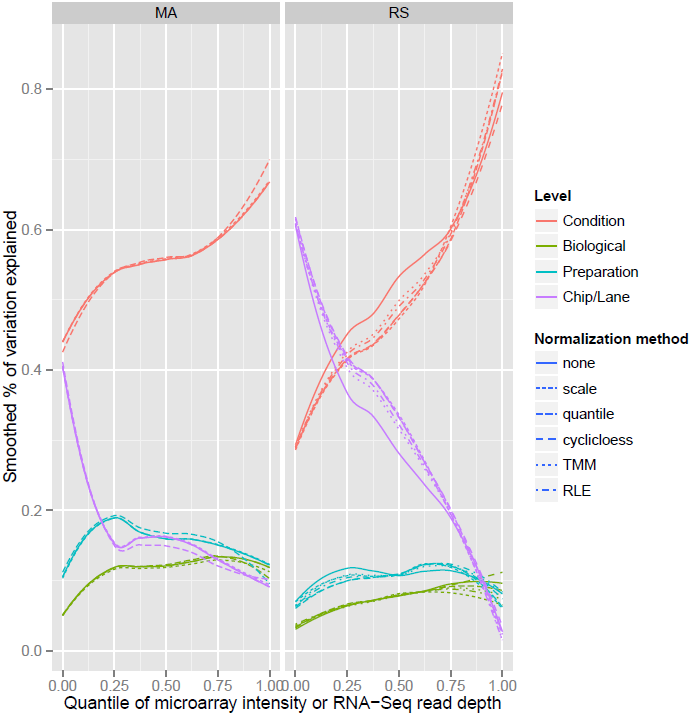
Percent of variance explained by each nested level of the experiment, comparing several normalization methods. “None”, “scale”, “quantile” and “cyclicloess” denote options used in `normalizeBetweenArrays` in the limma R package, TMM denotes the trimmed mean of M-values method of Robinson and Oshlack 2010 [34], and RLE denotes the “relative log expression” method of Anders and Huber 2010 [35]. TMM and RLE were applied only to the RNA-Seq data.

**Figure S4:**
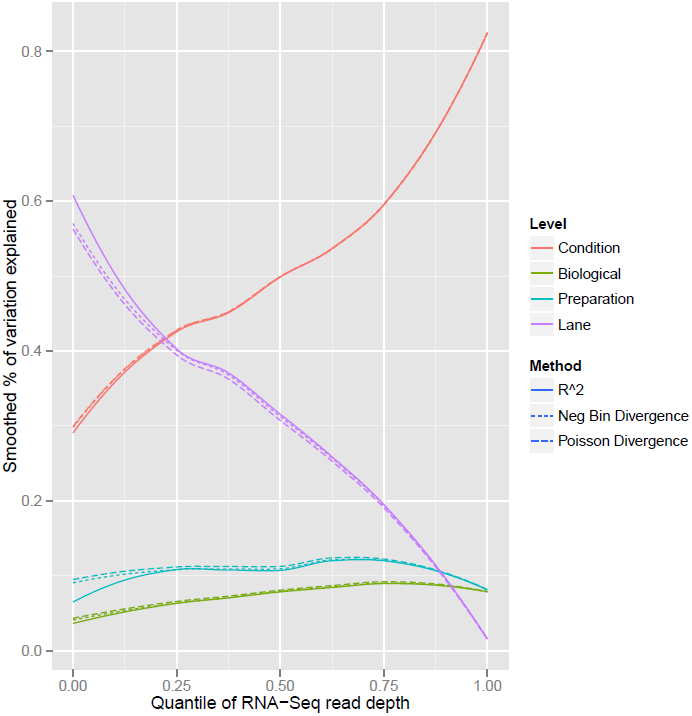
Comparison of an *R*^2^ from ANOVA computed for transformed RNA-Seq, using TMM normalization, to a pseudo-*R*^2^ metric making use of either a Poisson or a negative binomial model (**Materials and Methods**).

**Figure S5:**
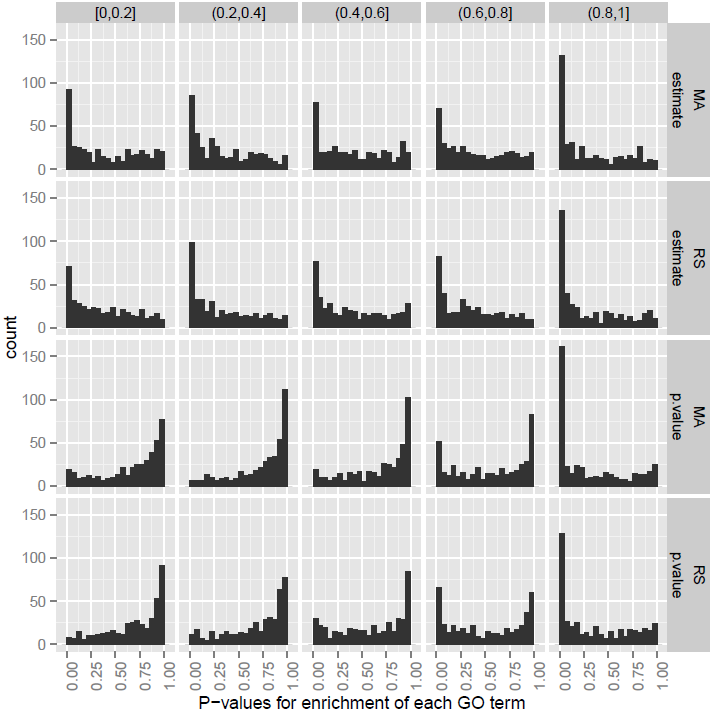
P-value histograms for Wilcoxon tests of gene set enrichment. The plot is divided vertically based on the technology (micrarray or RNA-Seq) and metric (log fold change estimate or p-value) used in the enrichment analysis, and horizontally based on the quantile of the average intensity within each gene.

**Figure S6:**
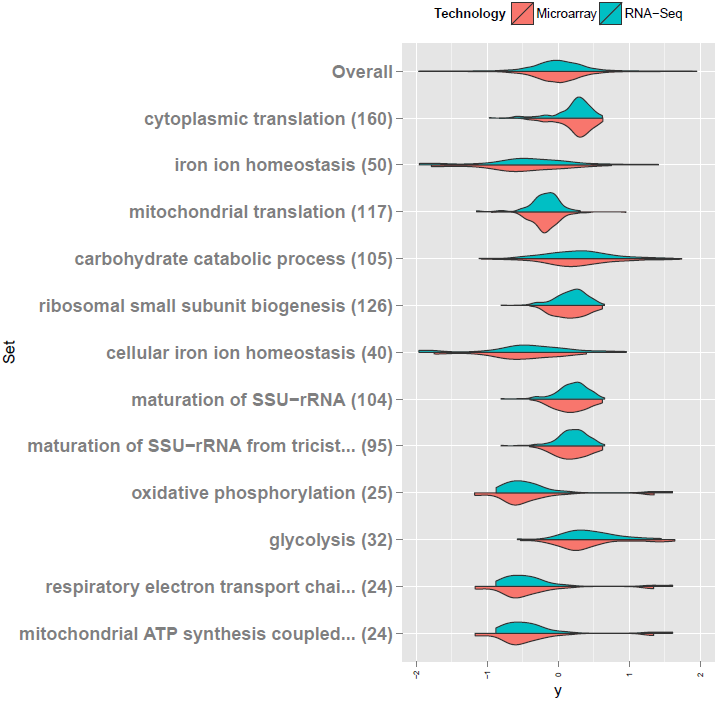
The 12 sets that were found most significantly differentially expressed in microarrays and RNA-Seq, within the Biological Process ontology.

**Figure S7:**
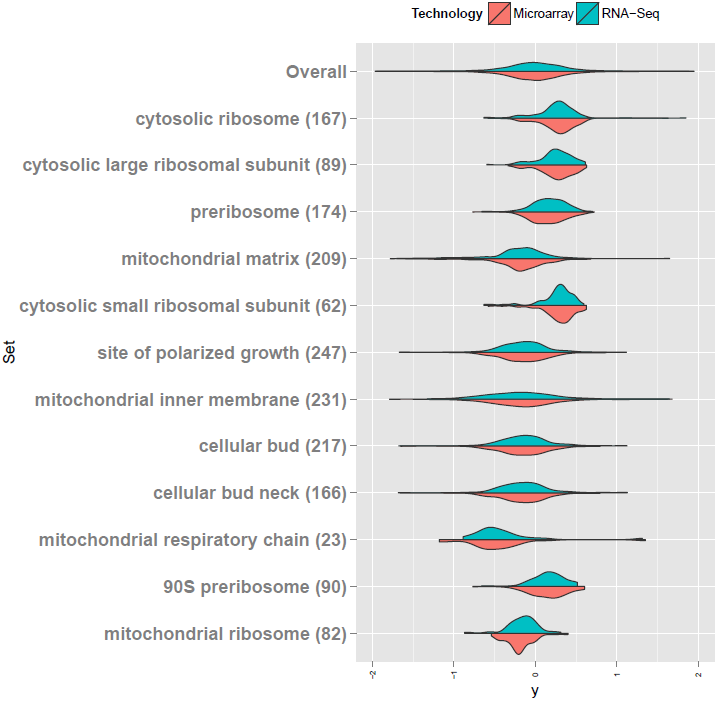
The 12 sets that were found most significantly differentially expressed in microarrays and RNA-Seq, within the Cellular Compartment ontology.

**Figure S8:**
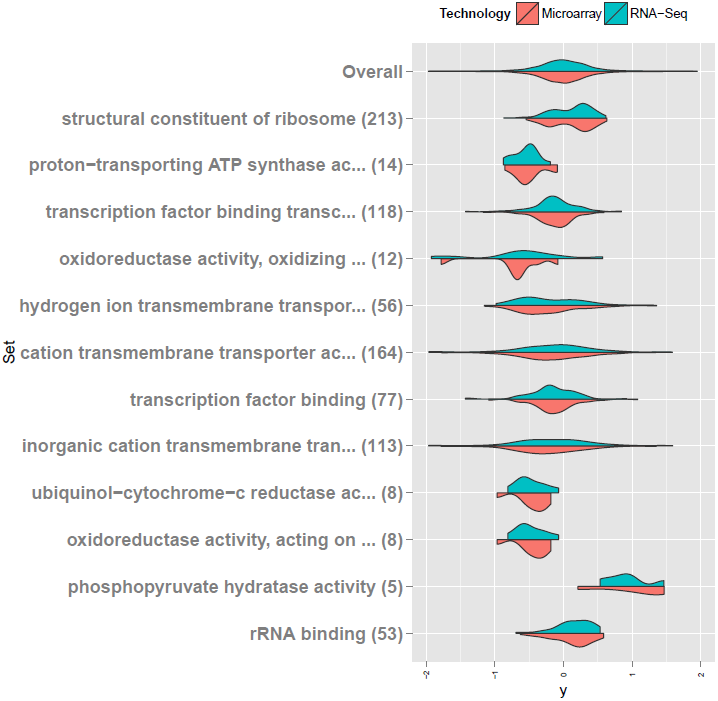
The 12 sets that were found most significantly differentially expressed in microarrays and RNA-Seq, within the Molecular Function ontology.

**Figure S9:**
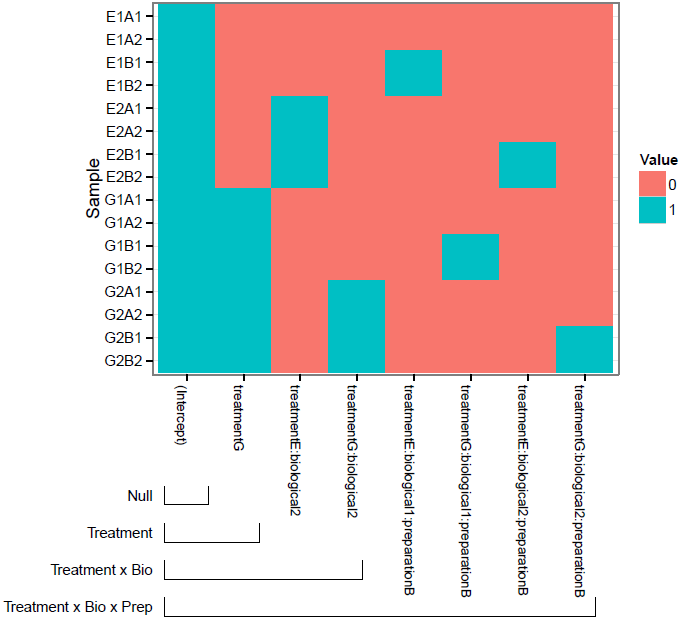
The design matrix *X* used to fit linear models to the nested experiment and determine *R*^2^. The Null, Treatment, Treatment×Biological, and Treatment×Biological×Preparation models use 1, 2, 4, or 8 columns of this matrix, respectively, as shown.

